# Machine Learning Made Easy (MLme): A Comprehensive Toolkit for Machine Learning-Driven Data Analysis

**DOI:** 10.1101/2023.07.04.546825

**Authors:** Akshay Akshay, Mitali Katoch, Navid Shekarchizadeh, Masoud Abedi, Ankush Sharma, Fiona C. Burkhard, Rosalyn M. Adam, Katia Monastyrskaya, Ali Hashemi Gheinani

## Abstract

**Background:** Machine learning (ML) has emerged as a vital asset for researchers to analyze and extract valuable information from complex datasets. However, developing an effective and robust ML pipeline can present a real challenge, demanding considerable time and effort, thereby impeding research progress. Existing tools in this landscape require a profound understanding of ML principles and programming skills. Furthermore, users are required to engage in the comprehensive configuration of their ML pipeline to obtain optimal performance.

**Results:** To address these challenges, we have developed a novel tool called *Machine Learning Made Easy* (MLme) that streamlines the use of ML in research, specifically focusing on classification problems at present. By integrating four essential functionalities, namely Data Exploration, AutoML, CustomML, and Visualization, MLme fulfills the diverse requirements of researchers while eliminating the need for extensive coding efforts. To demonstrate the applicability of MLme, we conducted rigorous testing on six distinct datasets, each presenting unique characteristics and challenges. Our results consistently showed promising performance across different datasets, reaffirming the versatility and effectiveness of the tool. Additionally, by utilizing MLme’s feature selection functionality, we successfully identified significant markers for CD8+ naive (BACH2), CD16+ (CD16), and CD14+ (VCAN) cell populations.

**Conclusion:** MLme serves as a valuable resource for leveraging machine learning (ML) to facilitate insightful data analysis and enhance research outcomes, while alleviating concerns related to complex coding scripts. The source code and a detailed tutorial for MLme are available at https://github.com/FunctionalUrology/MLme.

**Key Points:**

- MLme is a novel tool that simplifies machine learning (ML) for researchers by integrating Data Exploration, AutoML, CustomML, and Visualization functionalities.
- MLme improves efficiency and productivity by streamlining the ML workflow and eliminating the need for extensive coding efforts.
- Rigorous testing on diverse datasets demonstrates MLme’s promising performance in classification problems.
- MLme provides intuitive interfaces for data exploration, automated ML, customizable ML pipelines, and result visualization.
- Future developments aim to expand MLme’s capabilities to include support for unsupervised learning, regression, hyperparameter tuning, and integration of user-defined algorithms.

## Introduction

In the realm of research, machine learning (ML) has emerged as a vital resource for analyzing intricate datasets that conventional statistical approaches struggle to interpret^1–5^. However, the integration of machine learning (ML) into research presents a multitude of challenges. Foremost, the construction and execution of an effective ML pipeline can be daunting, requiring deep domain expertise, extensive technical knowledge, and proficient programming skills. In addition, the utilization of ML techniques demands a comprehensive understanding of the underlying principles to ensure that the trained models are unbiased and transparent.

Multiple tools have been developed to streamline the process of building and executing ML pipelines (Table S1)^6–9^. These tools often require a significant level of coding proficiency and extensive configuration to achieve optimal effectiveness. Additionally, many of these tools, serve as algorithm recommenders, functioning by running multiple ML algorithms on user-provided data and providing model performance metrics. However, this approach can limit user input and guidance, as the tools tend to prioritize automated decision-making rather than allowing users to actively participate in the process. As a result, tailoring the ML models to specific research needs and ensuring that the models align with domain knowledge and expertise can be challenging. This lack of flexibility and limited user control potentially hinders the accuracy and applicability of the research outcomes.

*Machine Learning Made Easy* (MLme) is a comprehensive solution aimed at bridging the gap between researchers and the inherent technical complexities of ML. It facilitates the adoption of ML techniques by simplifying the ML workflow and minimizing the typically steep learning curve associated with ML. Through its intuitive interfaces, MLme enhances accessibility and usability for researchers of varying levels of technical expertise (Figure 1).

**Figure 1.**
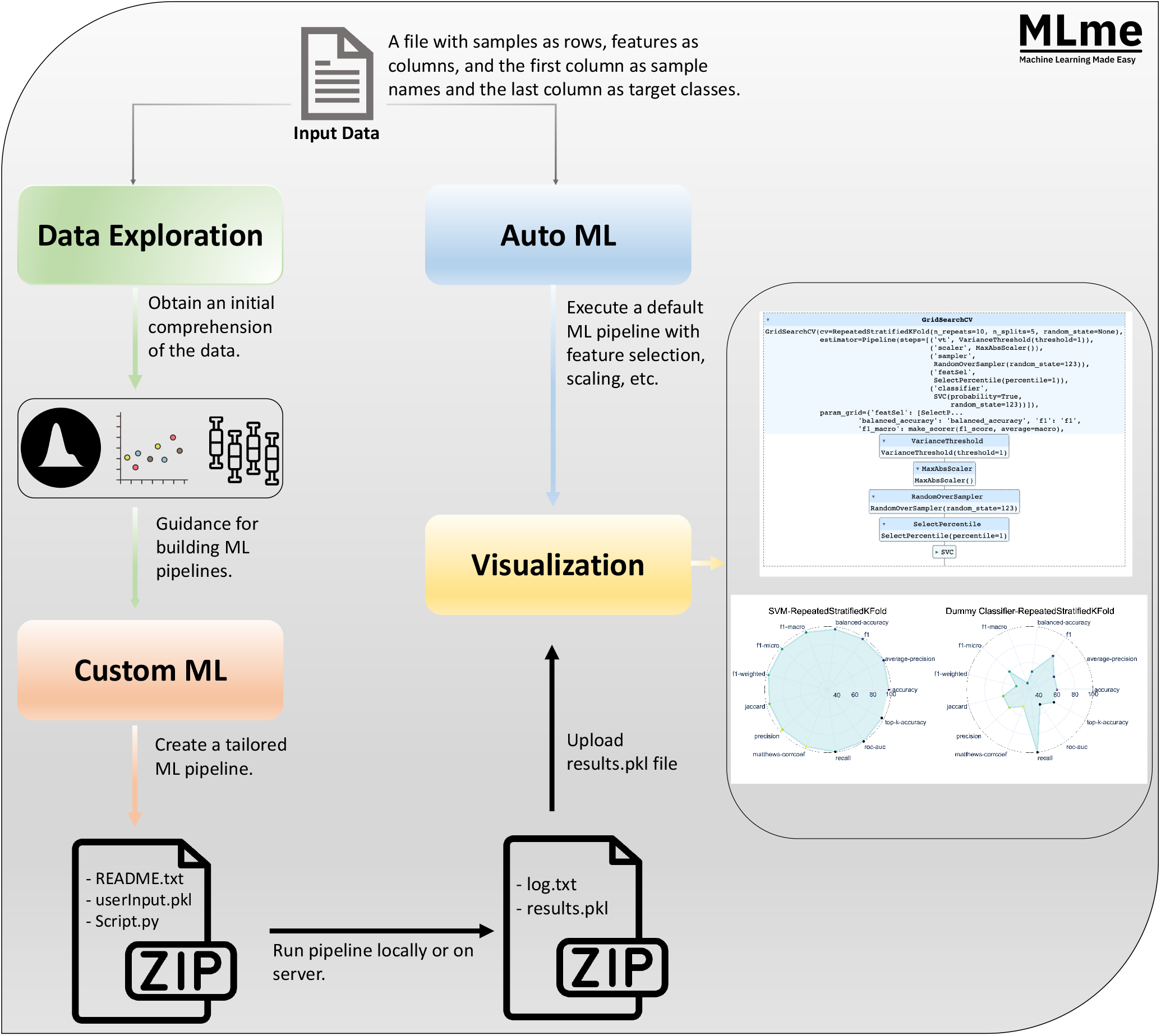
Graphical abstract. The input data for Machine Learning Made Easy (MLme) is a file with samples as rows and features as columns, with sample names in the first column and target classes in the last column. MLme provides various features to enhance usability. The data exploration feature enables users to explore the data and gain initial insights. For advanced users, the custom ML feature allows the creation of custom ML pipelines. Upon execution, MLme generates a compressed zip file containing inputParameter.pkl, script.py, and README.txt. Alternatively, users can opt for the AutoML feature, which applies a default ML pipeline to the input file. Both customML and AutoML produce a results.pkl file, which can be further analyzed using the visualization feature.

MLme offers four important components: Data Exploration, AutoML, CustomML, and Visualization, each serving a specific purpose in understanding and extracting meaningful information from the data within the ML workflow. Through the intuitive Data Exploration feature, users easily examine their datasets and gain preliminary understanding using an interactive interface. For advanced users, the CustomML interface within MLme provides a flexible platform to design and develop tailor-made ML pipelines that align with their specific research requirements. Furthermore, it facilitates effortless interpretation and analysis of results with rich visualization capabilities.

## Key features of *Machine Learning Made Easy* (MLme)

MLme is a multifaceted toolkit that equips researchers with the functionalities necessary to effectively utilize ML in their research. It consists of four distinct web interfaces, each tailored to address specific research needs, ensuring a versatile and comprehensive experience for users.

### Data Exploration

The Data Exploration feature of MLme allows users to upload their datasets and explore them using a range of statistical visualizations, such as density plots, scatter matrix plots, area plots, and class distribution plots (Figure S1.A). These visualizations and statistical summaries enable users to gain a comprehensive understanding of their data, including patterns and trends within the data, data distribution, and potential outliers. A density plot, for instance, can reveal how data is distributed, while a scatter matrix plot can identify potential correlations. Class distribution plots are particularly useful for comprehending the balance of target classes within the dataset, which can be crucial when designing a machine learning model.

Overall, the Data Exploration feature enables users to efficiently explore their datasets and acquire initial insights into their data. This knowledge can inform subsequent modeling decisions, ensuring that users are using the most appropriate modeling techniques for their specific dataset.

### AutoML

The AutoML feature in MLme enables users to effortlessly extract meaningful information from their datasets using ML, even without extensive technical expertise (Figure S1.B). With a preconfigured ML pipeline (Figure 2), the AutoML handles essential preprocessing steps such as data resampling, scaling, and feature selection. These steps ensure that the input data is properly prepared for ML algorithms, enhancing the performance and reliability of subsequent trained models. The AutoML conducts training and evaluation of multiple classification models, including a dummy classifier. By employing diverse models, users gain a comprehensive understanding of their data and can identify the most effective algorithms for their specific dataset.

**Figure 2.**
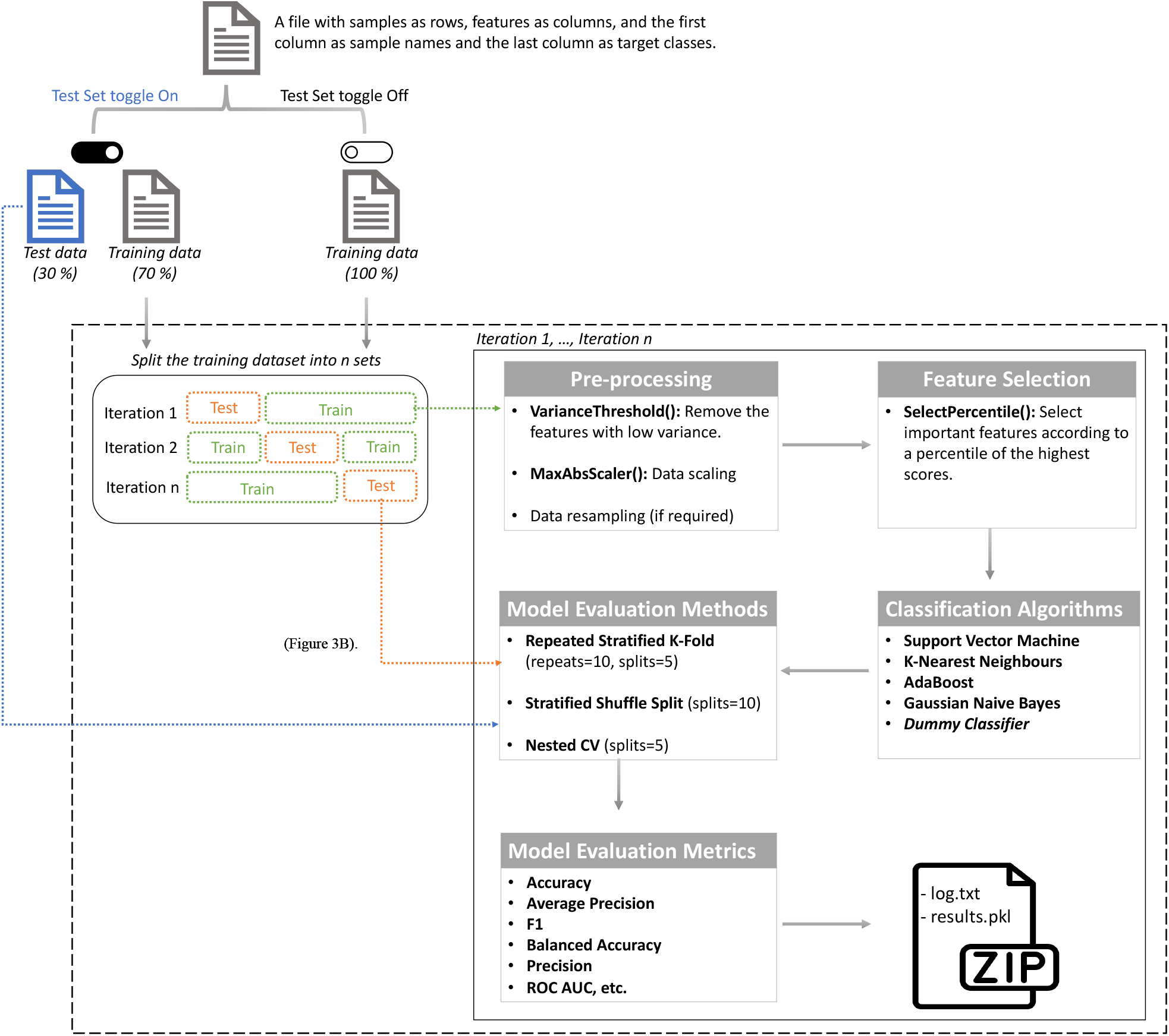
Default ML Pipeline for AutoML. The default ML pipeline can be represented as a flowchart that starts by splitting the input dataset into training and independent test sets, provided the user has activated the test set option. Otherwise, the entire dataset is used for training. In the subsequent step, the training dataset is divided into n bins of equal size through stratified sampling. From these bins, k-1 are designated as training sets while the remainder becomes the test set. In the pre-processing step, low variance features are removed first, followed by data scaling and resampling. Subsequently, the SelectPercentile univariate feature selection method is applied to select important features, and five ML classification algorithms are trained. Model performance is assessed on the test set using three different methods, and multiple performance metrics are computed. This entire process is repeated for each unique bin in the k-fold CV method. The pipeline outputs a zip file comprising the log .txt and the results.pkl files. The user can examine the results by visualizing the contents of the pickle file using Machine Learning Made Easy (MLme).

After the pipeline is completed, the AutoML offers users various options for examining and interpreting the results. These options include intuitive and interactive plots, which help users gain a deeper understanding of the performance characteristics of the models. Additionally, users have the flexibility to download the results and explore them further using the Visualization interface at their convenience.

### CustomML

The CustomML feature of MLme empowers users with moderate to advanced knowledge of the ML domain to design and customize an ML pipeline that caters to their specific research needs (Figure S2.A). With its user-friendly and intuitive interface, users can easily include or exclude steps and algorithms using a simple toggle button. This eliminates the worry about writing complex programming scripts and allows to focus on selecting the most suitable steps and algorithms for the dataset.

CustomML offers an extensive range of preprocessing options, including seven algorithms for data resampling, nineteen algorithms for scaling, and a diverse array of feature selection algorithms to select relevant features from the dataset. Moreover, with sixteen classification algorithms available, users can refine their pipeline to align with their research requirements. To provide a comprehensive understanding of the trained model’s performance, CustomML supports ten different evaluation methods and fourteen evaluation metrics.

The customization options of CustomML are enhanced by allowing users to select the parameters value for all the provided algorithms, giving them greater control over the behavior of their developed pipeline. Once the pipeline is designed, it can be conveniently downloaded and executed either locally or on a cluster, offering flexibility in computing resources. The CustomML-generated ML pipeline produces a pickle file (.pkl) as an output upon completion, which contains all the results from the pipeline. This file can be uploaded to the Visualization interface, enabling users to interpret these results using various plots.

### Visualization

The Visualization feature in MLme allows users to effortlessly interpret their results without the need for advanced programming skills or expertise in data visualization (Figure S2.B). It provides a comprehensive range of plots and tables, covering fundamental as well as advanced options such as bar plots, heatmaps, and spider plots. These diverse visualization tools facilitate effective comparison of trained model performance.

Furthermore, this feature allows users to customize the appearance of their plots by selecting from over fifty different color palettes. Additionally, all generated plots are of high quality and are downloadable in high resolution, ensuring they are suitable for publication purposes.

## Use-Cases

### Dataset Selection Criteria

The MLme application is evaluated using six distinct datasets (Table S2) that are carefully chosen to ensure robustness. Factors such as sample size, diversity, class imbalance, and dimensionality are considered during the selection process. The selected datasets vary in sample size and diversity, providing a comprehensive assessment of the MLme application’s performance across different data scales.

This includes datasets of varying sizes, from small (chronic cymphocytic leukemia and cervical cancer study) to large (invasive breast carcinoma datasets), which test the application’s scalability and efficiency. Imbalanced datasets, like invasive Breast Carcinoma (BRCA), are included to evaluate the MLme application’s handling of class imbalance and prediction accuracy, which is particularly relevant in real-world scenarios, such as biological research. The datasets also address the challenge of high-dimensional features and low sample sizes, known as the curse of dimensionality. By including such datasets, the MLme ability to handle challenges is thoroughly assessed.

Furthermore, the Glass Identification dataset was selected as a non-biological example, offering variation, and enabling testing across diverse domains. This dataset, with multiple target classes, allows evaluation of the MLme application’s performance in multi-class classification problems.

### Dataset Descriptions

The first dataset comprised of mRNA patient data (n=136) obtained from a study on Chronic Lymphocytic Leukemia (CLL), which measured their transcriptome profiles^10^. Our objective was to build a model that could classify male and female patients based on their transcriptomic profiles, using the top 5,000 most variable mRNAs (excluding Y chromosome genes). The second dataset was collected from a cervical cancer study that analyzed the expression levels of 714 miRNAs in human samples (n=58)^11^.

The third and fourth datasets were obtained from The Cancer Genome Atlas (TCGA), consisting of mRNA (n=1219) and miRNA (n=1207) sequencing data from patients with invasive BRCA, which were retrieved using the TCGAbiolinks package^12^ in *R*. For the BRCA mRNA dataset, we focused only on differentially expressed genes from edgeR (FDR ≤ 0.001 and logFC > ±2)^13^. Our goal was to train a model capable of distinguishing normal and tumor samples for both cervical cancer and TCGA-BRCA datasets.

The fifth dataset consists of scRNA-seq data obtained from peripheral blood mononuclear cells (PBMCs) that were sequenced using 10× chromium technology^14^. Among all the cell populations described in this study, we specifically utilized the scRNA datasets of CD8+ naive, CD14+, and CD16+ monocytes (n=1500) with the goal of identifying distinct markers for each of these cell populations.

The sixth dataset utilized in this study was the widely recognized Glass Identification dataset (n=214) obtained from the University of California Irvine (UCI) ML repository^15^. This dataset comprises 10 distinct features that represents oxide content of glass samples. The primary objective of this dataset is to classify different types of glass based on their oxide content.

## Results

To perform a thorough assessment of the MLme functionality, we utilized its customML feature to construct distinct ML pipelines for CLL, Cervical cancer and TCGA datasets. These pipelines entailed various processing steps, including data scaling and resampling using different algorithms, multiple ML classifiers, diverse evaluation methods, and metrics. Additionally, we employed the AutoML feature of MLme to train multiple models for both the PBMC and glass datasets. The top-performing models consistently achieved scores exceeding 90% for all computed metrics across all evaluated datasets except Glass Identification dataset. As anticipated, the dummy classifiers performed the worst among all the datasets (Figures S3-S8).

To further demonstrate the applicability of MLme, we utilized its feature selection functionality from AutoML to identify the most important genes for classifying CD8+ naive, CD14+, and CD16+ monocyte cell populations from the PBMC dataset. By selecting the top 10% of the original input of 500 highly variable genes, MLme provided a list of 50 genes that are sufficient for classifying these cell types (Figure 3A). These 50 genes exhibited a strong correspondence with their respective cell populations, except for 13 ribosomal genes (RPS and RPL) that showed similar expression levels across all three cell types.

**Figure 3.**
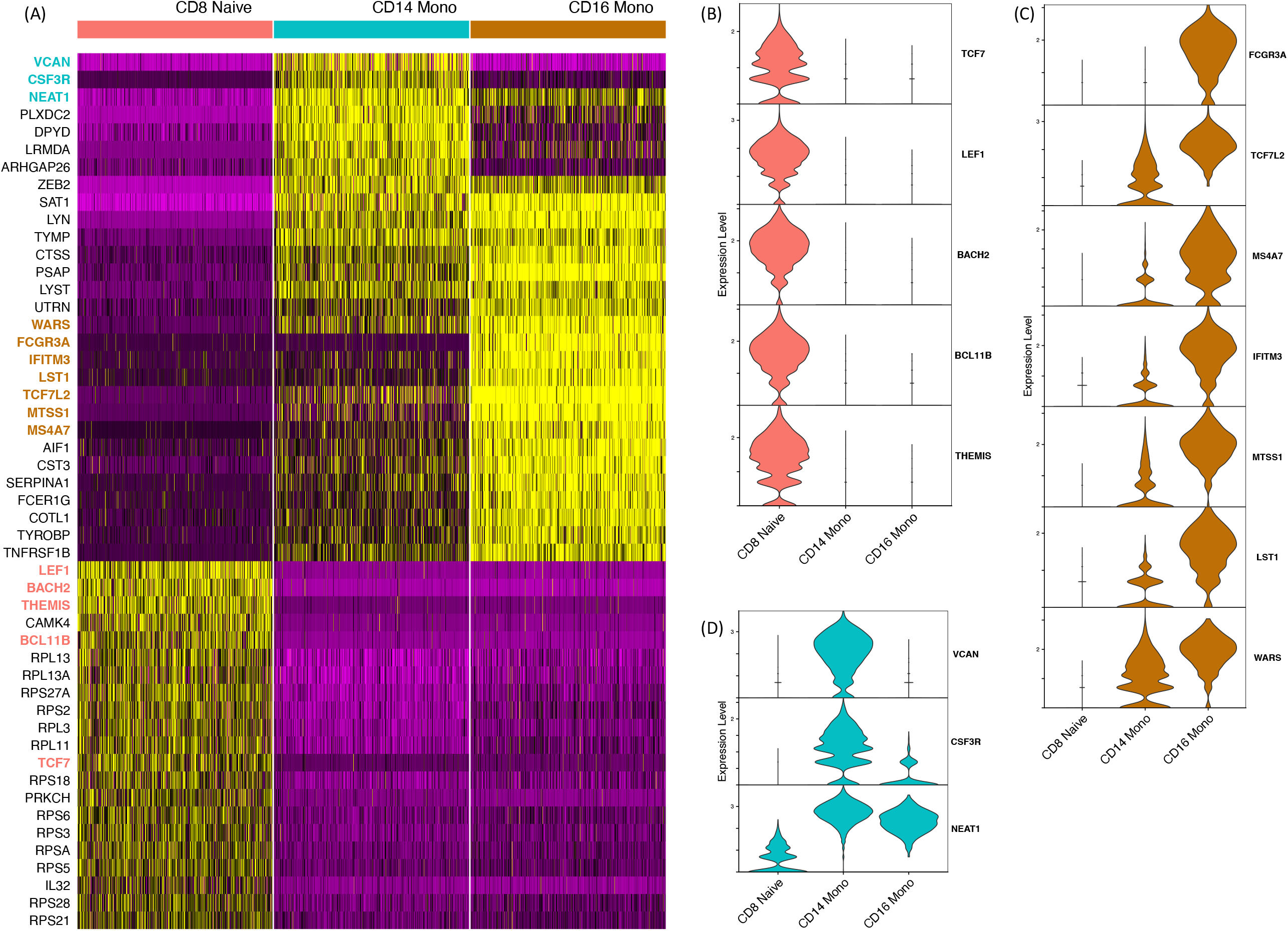
Identification of Potential Markers for CD8+ Naïve, CD16+, and CD14+ Cell Populations in PBMC Dataset. **(A)** Heatmap visualization showing the expression patterns of 50 genes selected by MLme. **(B), (C), and (D)** Demonstrate the expression levels of key markers specific to CD8+ naïve, CD16+, and CD14+ cell populations, respectively, within each cell type.

Among the remaining 37 genes, we discovered classic markers for CD8+ naive cells (TCF7^16,17^, LEF1^16^, BACH2^18^, BCL11B^19^, and THEMIS^20^), which have been previously described in the literature (Figure 3B). The list also included markers for the CD16+ cell population, such as FCGR3A (CD16), TCF7L2, MS4A7, IFITM3, MTSS1, LST1, and WARS (Figure 3C), which have been associated with CD16+ cells in previous studies^21,22^. Furthermore, our marker list encompassed known CD14+ specific genes, including VCAN, a marker of monocytic lineage^23^, CSF3R, previously described in the CD14+ population^24^, and NEAT1 (Figure 3D). These findings validate the biological relevance of the selected genes and highlight the utility of the MLme tool in biomedical research.

## Implementation

The MLme is developed using *Dash* library^25^ in the *Python*^26^ programming language. Plots are generated using *Plotly*^27^, *matplotlib*^28^, and *bokeh*^29^ libraries. *Pandas*^30^ and *NumPy*^31^ libraries are used to handle data storage and processing. The development of the ML pipeline is facilitated by employing the *Scikit-Learn*^32^ and *Imbalanced Learn*^33^ libraries.

## Limitations

Currently, MLme focuses on classification problems since a substantial portion of research questions and available datasets are aligned with the domain of classification. This limitation hinders MLme applicability to regression or unsupervised learning tasks. Additionally, the tool lacks built-in hyperparameter tuning capabilities. This absence of a key feature may hinder users in fine-tuning their models.

Overall, while the current version of the MLme has limitations concerning regression and unsupervised learning problems, it excels in its core objective of addressing classification tasks. It is worth noting that users have the flexibility to choose values for all the parameters of a given algorithm through the user interface, to some extent mitigating the impact of the lack of built-in hyperparameter tuning to some extent.

## Conclusion

Our paper introduces a user-friendly tool called MLme, which offers a wide range of functionalities for ML analysis. Its primary goal is to make machine learning (ML) accessible to users of all skill levels by removing technical barriers. With the Data Exploration feature, users can efficiently explore datasets and gain initial insights into their data. The AutoML feature simplifies ML usage, allowing them to leverage ML capabilities without dealing with complex technicalities. Moreover, the CustomML functionality assists in creating personalized pipelines using an intuitive graphical user interface that caters to specific requirements, eliminating the need for coding complexities. Additionally, the visualization features enable users to interactively explore and understand model performance, without extensive data visualization or coding expertise. In summary, MLme is a powerful and user-friendly tool that empowers researchers to enhance their research outcomes through ML.

## Outlook

Despite the limitations mentioned above, there are several promising directions for future development of the MLme. Our primary objective is to expand the capabilities of MLme to include support for unsupervised learning and regression problems. This expansion will greatly enhance the tool’s utility and enable its application in a broader range of ML tasks.

Recognizing the importance of hyperparameter tuning in optimizing models, we plan to incorporate hyperparameter tuning capabilities into the tool. This addition will enable users to fine-tune their models and improve overall performance, thereby increasing the MLme effectiveness and reliability. Additionally, we intend to introduce a feature that allows users to upload and integrate their own algorithms into the pipeline. This feature will enable users to use their preferred algorithms, even if they are not currently available within the tool, thereby expanding its applicability and customization options.

These future developments aim to overcome the current limitations of MLme and enhance its functionality and adaptability. By addressing these limitations, we firmly believe that the MLme will evolve into a more comprehensive and valuable resource for ML practitioners.

## Availability of supporting source code and requirements

Project name: Machine Learning Made Easy (MLme)

Project home page: https://github.com/FunctionalUrology/MLme

Operating system(s): Platform independent

Programming language: Python (version 3.9)

Other requirements: Docker or Python

License: GNU GPL

## Author contributions statement

K.M., A.H.G, A.A., and M.K. conceived the idea for the manuscript. A.A. and M.K. wrote the source code and conducted testing and debugging of the MLme. K.M., F.C.B, R.M.A, A.H.G, and A.S. provided feedback on the biological application of the tool. N.S., M.A., A.H.G, and A.S. provided technical feedback throughout the development phase and participated in testing and debugging. K.M., A.S. and M.K. wrote the manuscript with inputs from all the other authors. All authors contributed to proofreading and revising the manuscript.

## Funding

We gratefully acknowledge the financial support of the Swiss National Science Foundation (SNF Grant 310030_175773 to FCB and KM, 212298 to FCB and AHG) and the Wings for Life Spinal Cord Research Foundation (WFL-AT-06/19 to KM). AHG and RMA are supported by R01 DK127673. MK is supported by the Else Kröner-Fresenius-Stiftung (EKFS 2021_EKeA.33). The authors acknowledge the financial support from the Federal Ministry of Education and Research of Germany and by the Sächsische Staatsministerium für Wissenschaft Kultur und Tourismus in the program Center of Excellence for AI-research “Center for Scalable Data Analytics and Artificial Intelligence Dresden/Leipzig” (project identification number: ScaDS.AI).

## Conflict of Interest

The authors have declared no competing interests.

## Data Availability

All supporting data, including the input dataset, ‘inputParameters.pkl’, and ‘results.pkl’ files, for all evaluated datasets, is available on Zenodo^34^. The ‘results.pkl’ files can be visualized using the Visualization feature of MLme. DOME-ML (Data, Optimisation, Model, and Evaluation in Machine Learning) annotation, supporting the current study, is available through DOME Wizard.

## Acknowledgment

We would like to express our sincere gratitude to Pedro Perreira Amado for his invaluable contribution in testing MLme.

## Notes

https://github.com/FunctionalUrology/MLme

