## Supplementary figure for "Machine Learning Made Easy (MLme): A Comprehensive Toolkit for Machine Learning-Driven Data Analysis"

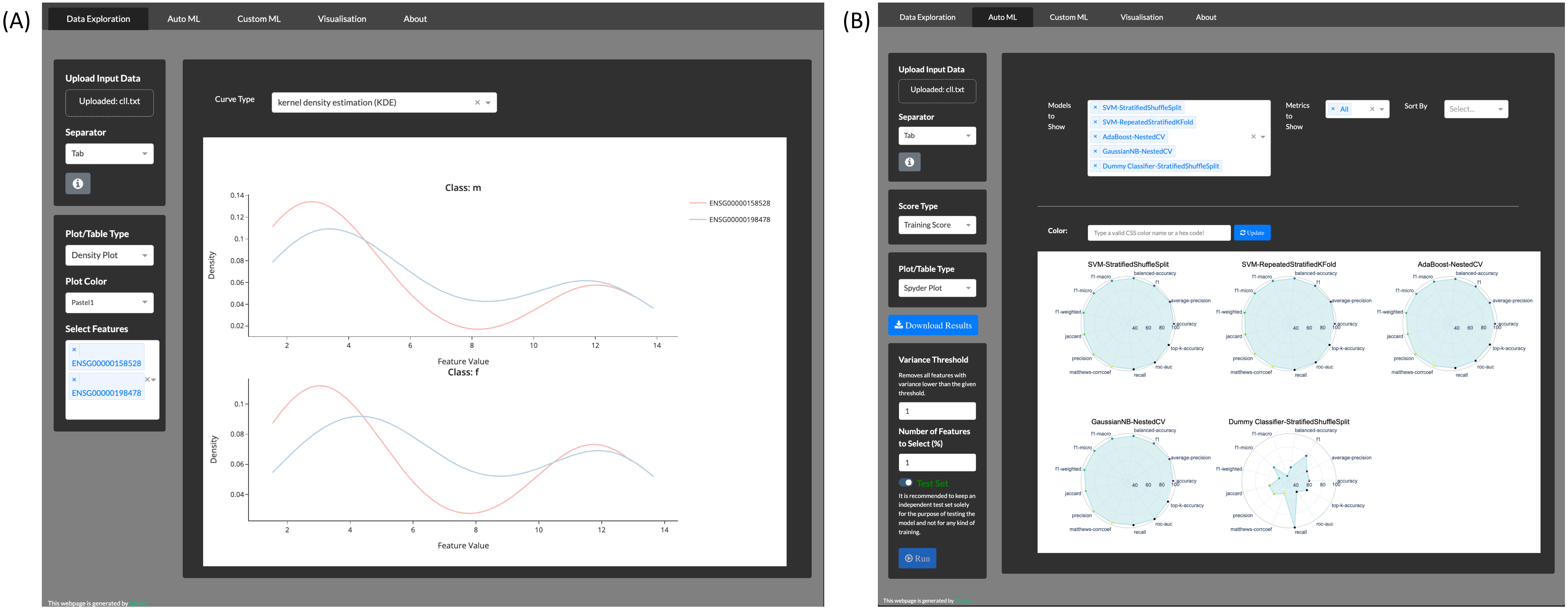


**Figure S1. Key features of Machine Learning Made Easy (MLme). (A)** Data Exploration, **(B)** AutoML.

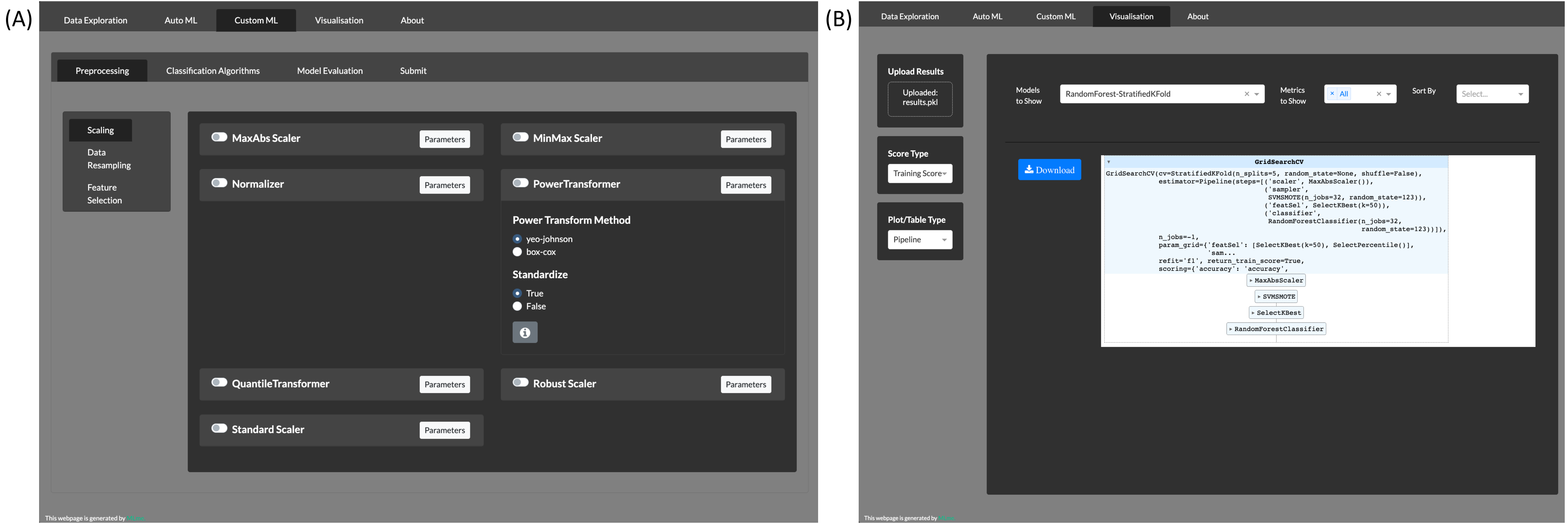


**Figure S2. Key features of Machine Learning Made Easy (MLme). (A)** CustomML, **(B)** Visualization.

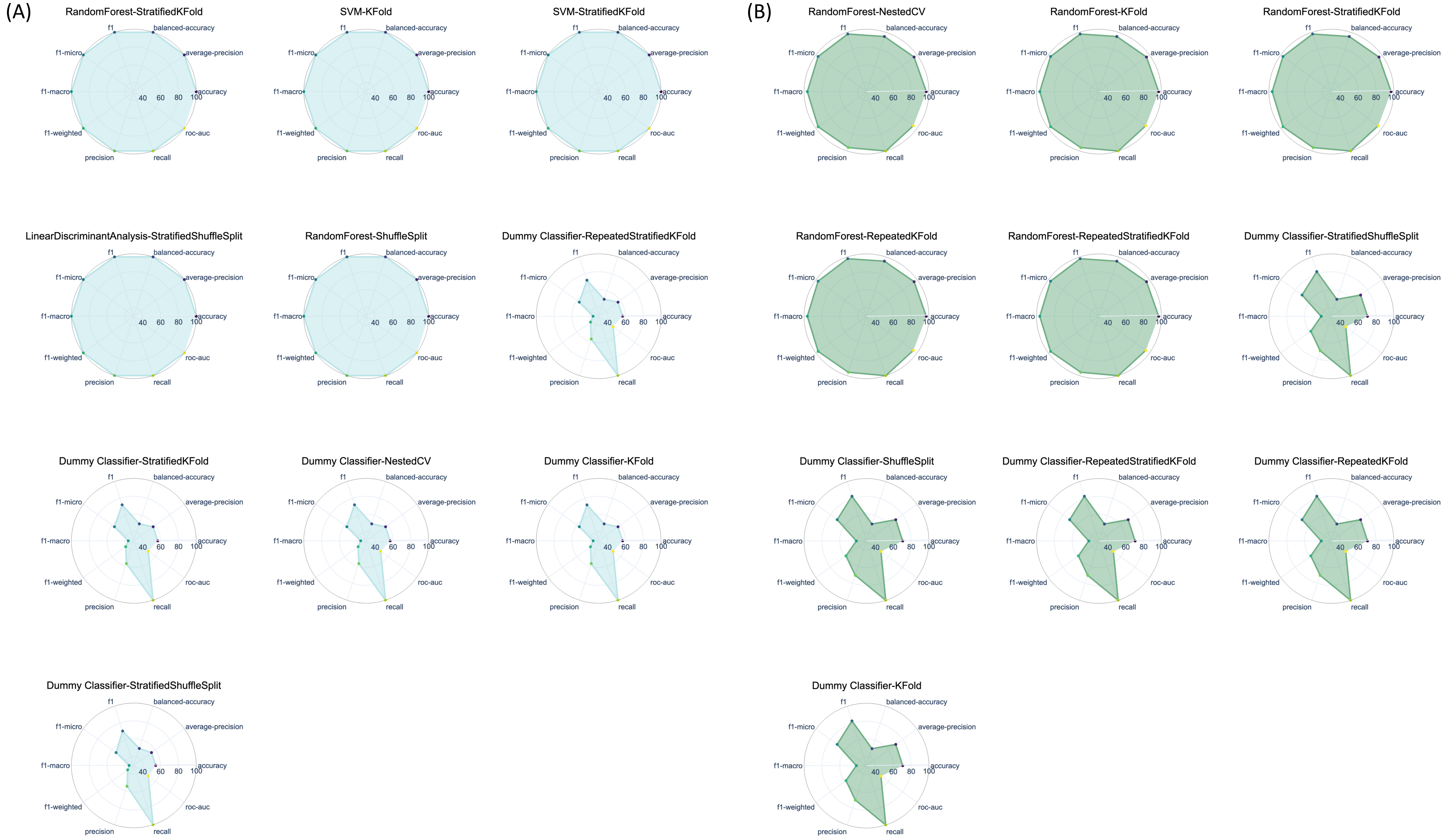


**Figure S3. Projection of metrics scores on two-dimensional (2D) polar coordinates.** The plots illustrate the performance scores of the top and worst five machine learning (ML) algorithms trained on the Chronic Lymphocytic Leukemia (CLL) dataset, both during training (A) and testing (B). Each ML model is represented by a circle, and each vertex represents a specific performance metric. A circle with a larger shaded area indicates better performance.

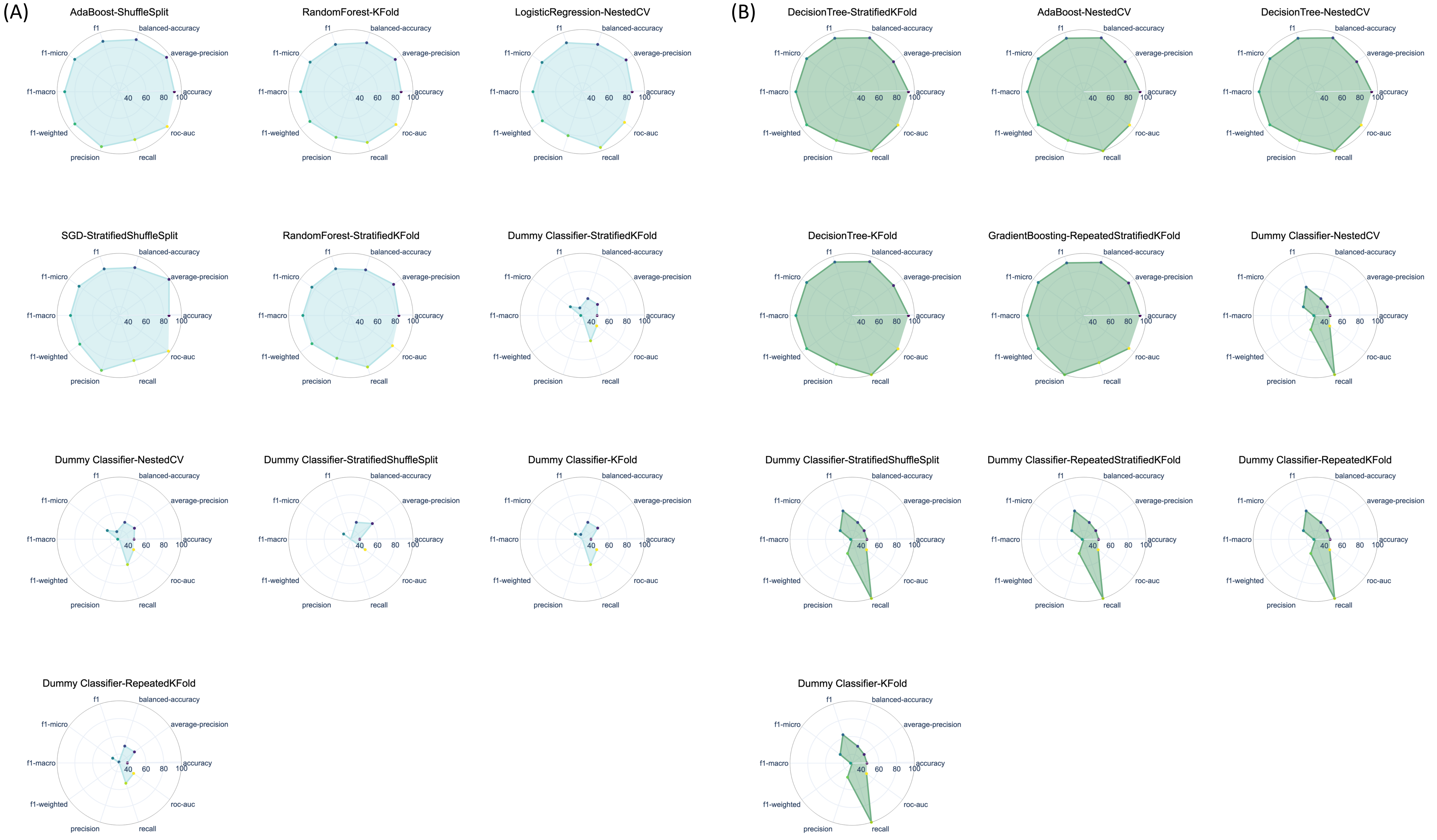


**Figure S4. Projection of metrics scores on two-dimensional (2D) polar coordinates.** The plots illustrate the performance scores of the top and worst five machine learning (ML) algorithms trained on the Cervical Cancer dataset, both during training (A) and testing (B). Each ML model is represented by a circle, and each vertex represents a specific performance metric. A circle with a larger shaded area indicates better performance.

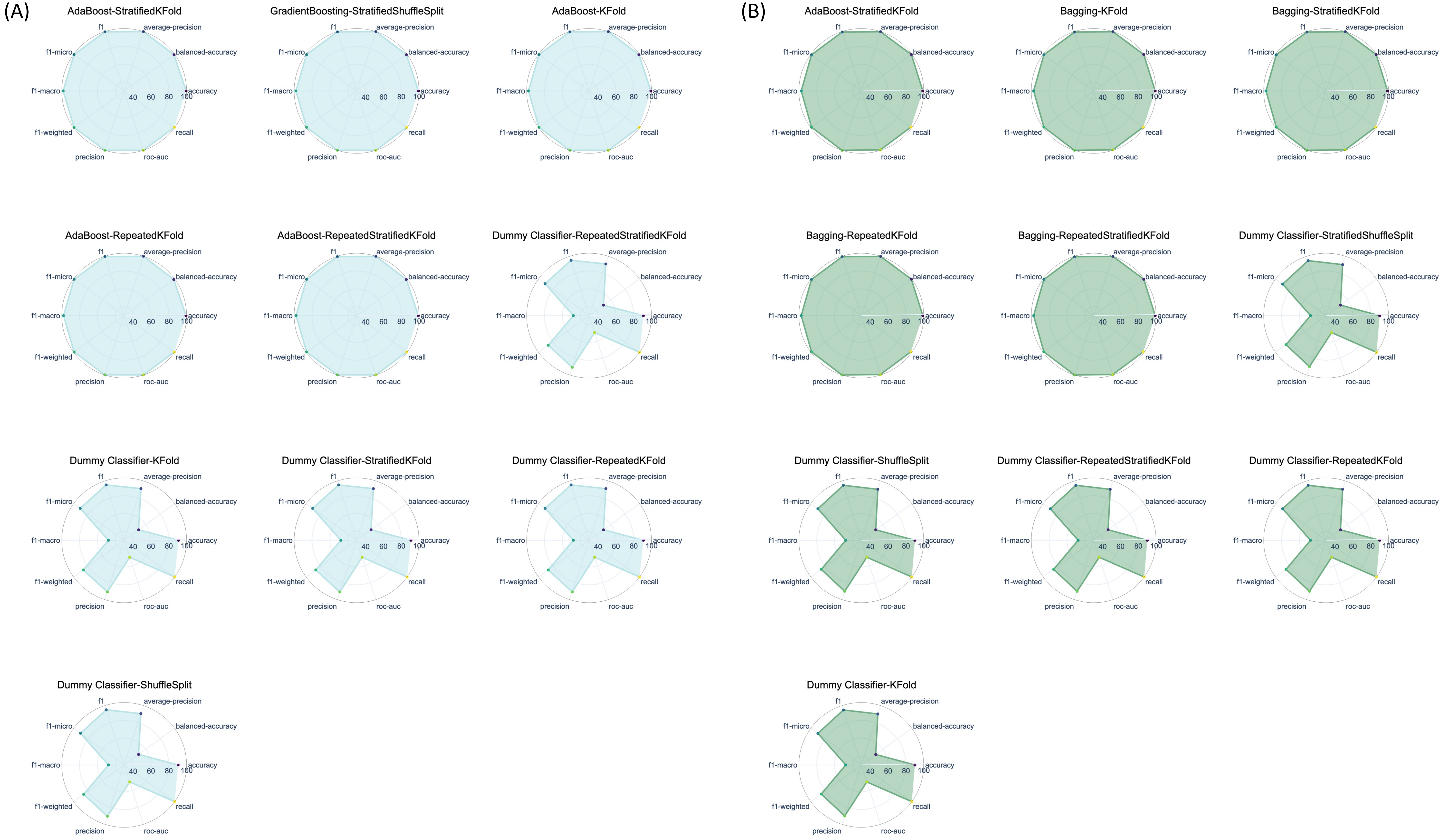


**Figure S5. Projection of metrics scores on two-dimensional (2D) polar coordinates.** The plots illustrate the performance scores of the top and worst five machine learning (ML) algorithms trained on the TCGA mRNA dataset, both during training (A) and testing (B). Each ML model is represented by a circle, and each vertex represents a specific performance metric. A circle with a larger shaded area indicates better performance.

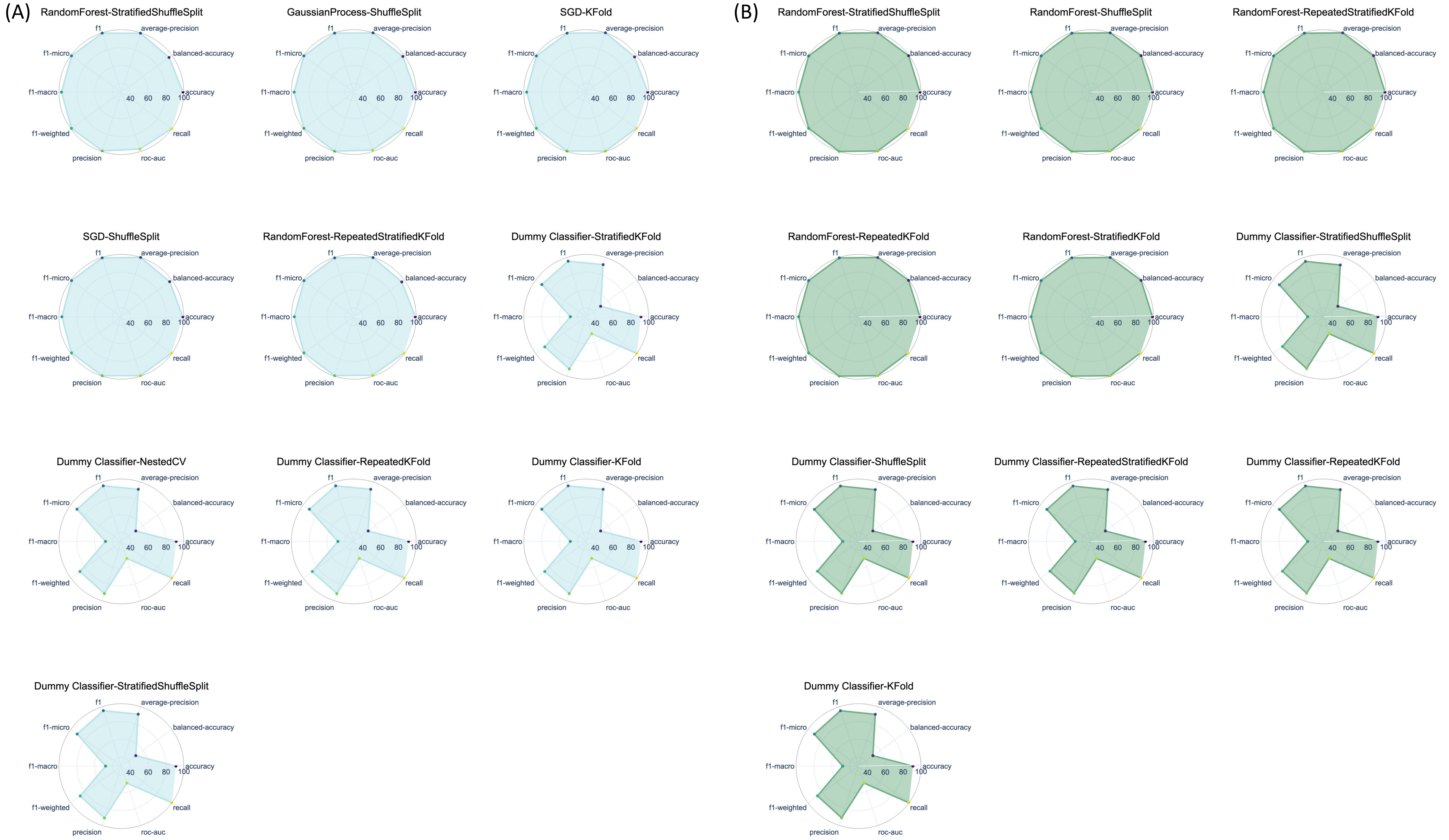


**Figure S6. Projection of metrics scores on two-dimensional (2D) polar coordinates.** The plots illustrate the performance scores of the top and worst five machine learning (ML) algorithms trained on the TCGA miRNA dataset, both during training (A) and testing (B). Each ML model is represented by a circle, and each vertex represents a specific performance metric. A circle with a larger shaded area indicates better performance.

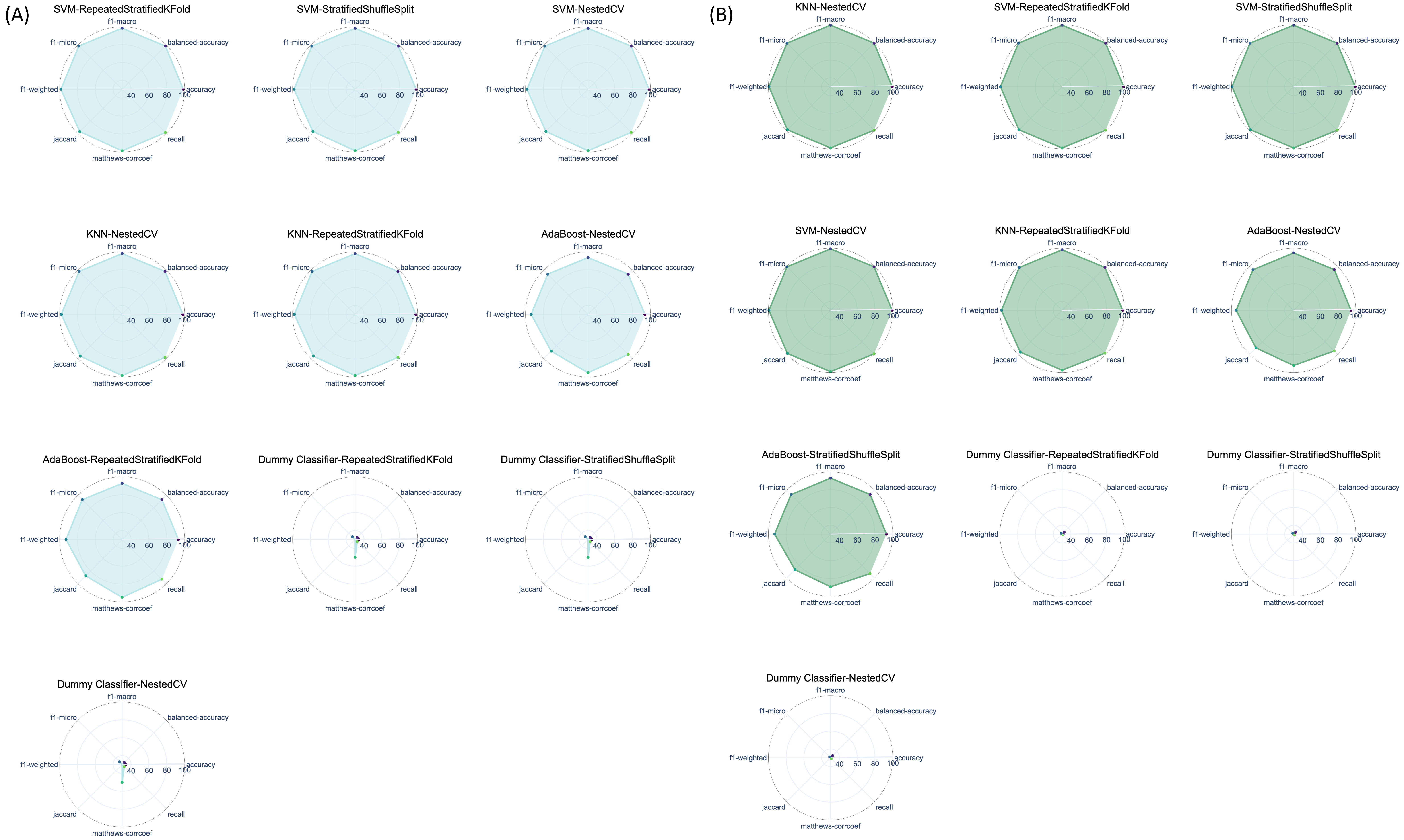


**Figure S7. Projection of metrics scores on two-dimensional (2D) polar coordinates.** The plots illustrate the performance scores of the top and worst five machine learning (ML) algorithms trained on the Peripheral Blood Mononuclear Cells (PBMC) dataset, both during training (A) and testing (B). Each ML model is represented by a circle, and each vertex represents a specific performance metric. A circle with a larger shaded area indicates better performance.

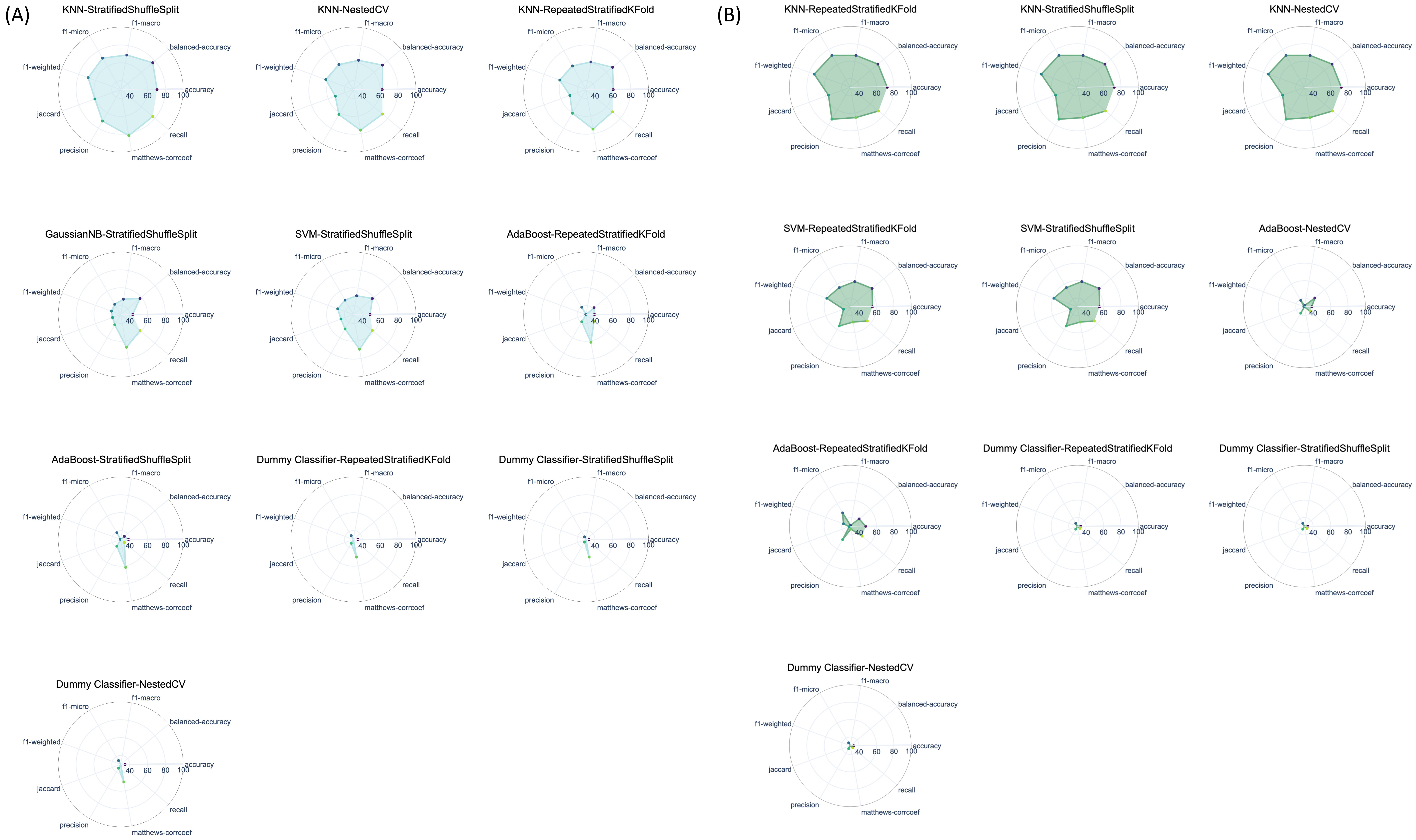


**Figure S8. Projection of metrics scores on two-dimensional (2D) polar coordinates.** The plots illustrate the performance scores of the top and worst five machine learning (ML) algorithms trained on the Glass Identification dataset, both during training (A) and testing (B). Each ML model is represented by a circle, and each vertex represents a specific performance metric. A circle with a larger shaded area indicates better performance.
