## Supplementary tables for "Machine Learning Made Easy (MLme): A Comprehensive Toolkit for Machine Learning-Driven Data Analysis"

| ***Tool*** | ***GUI*** | ***Data Exploration*** | ***Interactive Visualization*** | ***Design Custom Pipeline*** | ***Preprocessing*** |
|---|---|---|---|---|---|
| **MLme** | ✓ | ✓ | ✓ | ✓ | Scaling, Data Resampling, Feature Selection |
| TPOT | | | | ✓* | Feature Selection |
| PennAI | ✓ | | | | Feature Selection |
| AutoSklearn 2.0 | | | | ✓* | Scaling, Imputation, Feature Selection |
| HyperoptSklearn | | | | ✓* | Scaling |

*Table S1: Comparison of features between MLme and other similar machine learning automation tools. * = Coding expertise is required. GUI = Graphical User Interface.*

| ***Dataset*** | ***Data type*** | ***Number of Samples*** | ***Number of Features*** | ***Target Class ratio*** |
|---|---|---|---|---|
| CLL | mRNA | 136 | 5000 | Male (n=82): Female (n=54) |
| Cervical cancer | miRNA | 58 | 714 | Normal (n=29): Tumor (n=29) |
| TCGA-BRCA | miRNA | 1207 | 1404 | Normal (n=104): Tumor (n=1104) |
| TCGA-BRCA | mRNA | 1219 | 5520 | Normal (n=113): Tumor (n=1106) |
| PBMC | scRNA-seq | 1500 | 500 | CD8 Naive (n=500 cells) : CD14 Monocytes (n=500 cells) : CD16 Monocytes (n=500 cells) |
| Glass Identification | Oxide content (i.e., Na, Fe, K, etc) | 214 | 10 | Glass 1 (70), Glass 2 (76), Glass 3 (17), Glass 5 (12), Glass 6 (10), Glass 7 (29) |

*Table S2: Example datasets used in this study. CLL = Chronic Lymphocytic Leukemia. TCGA = The Cancer Genome Atlas. BRCA = Invasive Breast Carcinoma. PBMC = Peripheral Blood Mononuclear Cells.*
